# Unraveling the population history of Indian Siddis

**DOI:** 10.1101/101857

**Authors:** Ranajit Das, Priyanka Upadhyai

**Affiliations:** Manipal Centre for Natural Sciences (MCNS), Manipal University, Manipal, Karnataka, India; Department of Medical Genetics, Kasturba Medical College, Manipal University, Manipal, Karnataka, India

**Keywords:** Siddi, Geo-location, GPS, Admixture, Portuguese slave trade, Bantu expansion

## Abstract

The Siddis are a unique Indian tribe of African, South Asian and European ancestry. While their ancestral origins have been traced to the Bantu populations from sub-Saharan Africa, their population history has remained an enigmatic question. Here, we have traced the biogeographical origin of the Siddis employing an admixture based algorithm, Geographical Population Structure (GPS). We evaluated 14 Siddi genomes in reference to 5 African populations from the 1000 Genomes project and 7 Bantu populations from the Human Genome Diversity project. GPS assigned the Siddi genomes to west Zambia and the present-day border between Zimbabwe and northeastern Botswana, overlapping with one of the principal areas of secondary Bantu settlement in Africa, ~1700 years before present (YBP). This is concordant with the secondary Bantu dispersal route from the east African Bantu center that brought the African ancestors of the Siddis to settlement sites in southeast Africa, from where they were disseminated to India, by the Portuguese. Our results also suggest that while the Siddi genomes are significantly different from that of the Bantus, they displayed the highest genomic proximity to the Luhyas and North-East Bantus from Kenya, and that ancestral Siddis are likely to have split from the Luhyas, ~2700 YBP, in congruence with known Bantu expansion and population migration routes. Together with historical, linguistic and anthropological evidences our findings shine light on the genetic relatedness between populations, fine-scale population structure and recapitulate the population history of the Siddis, in the ethnohistorical context of India.

## Introduction

India and its adjoining areas in South Asia are a panorama of astounding ethnic, linguistic and genetic diversity interlaced with distinctive sociocultural practices. Contemporary India is a conglomeration of 4635 anthropologically well-defined populations, including 532 tribes (Narang, et al. 2010; Tamang, et al. 2012). It has witnessed numerous waves of immigration and gene-flow from various parts of the world, in prehistoric and historic times that have been instrumental in shaping its population structure and demography (Barnabas, et al. 1996; Kivisild, et al. 1999; Metspalu, et al. 2004; Misra 2001; Ratnagar 1995). The Siddis are a unique tribal group of African ancestry predominantly found in the Indian states of Gujarat, Karnataka, Andhra Pradesh and Telengana (Lodhi 1992). The earliest evidences of their migration that date back to ~1100 A.D indicate that the Siddis settled on the western coast of India (D. K 1969; Gauniyal, et al. 2008). By the 13^th^ century a greater influx of Siddis occurred and they were imported by regional Indian kings and princes to serve as slaves and soldiers (Shah, et al. 2011). Subsequently during the 16^th^-19^th^ centuries Siddis were transported in large numbers to India as slaves by the Portuguese (Bhattacharya 1970; Nevet 1981). Several previous investigations have suggested that the Siddi genomes are predominantly closest to that of the Africans (Thangaraj, et al. 1999). Further work not only confirmed the African ancestry of the Siddi people, but additionally revealed that they have assimilated considerable fractions of non-African admixture components in their genomes (Gauniyal, et al. 2011; Gauniyal, et al. 2008; Ramana, et al. 2001). This is not unexpected given that the Siddis shared long periods of contact with both the South Indians (South Asians) and the Portuguese (Europeans). In congruence, subsequent studies delineated three ancestral components, African, South Asian and European in the Siddi genomes (Gauniyal, et al. 2011; Gauniyal, et al. 2008; Shah, et al. 2011). Further analyses traced the ancestry of Siddis to the Bantu speaking populations from sub-Saharan Africa (Shah, et al. 2011). And admixture between the ancestral African Siddi immigrants and the South Indian populations was estimated to have likely occurred ~200 years before present (YBP) (Shah, et al. 2011). However, given the distinctive nature of Siddi ancestry their population history has remained relatively obscure, particularly in the ethnohistorical context of the genetically and culturally diverse Indian populace.

To facilitate our understanding of the population history of the Siddi tribe from India, we performed a genome-wide analysis of the Siddis (N=14), wherein samples had been procured from the Indian states of Gujarat (N=8) and Karnataka (N=6) by the Reich Lab at Harvard Medical School, USA (Moorjani, et al. 2013). An in-depth genetic comparison was carried out of the Siddis with the 5 African populations from the 1000 Genomes project, namely Yorubans from Ibadan, Nigeria (YRI) (N=108), Luhyas from Webuye, Kenya (LWK) (N=99), Gambians from the Western Divisions in Gambia (GWD) (N=113), Mendes from Sierra Leone (MSL) (N=85), and Esans from Nigeria (ESN) (N=99) (Genomes Project, et al. 2015). We further supplemented our analysis by evaluating the Siddis in reference to 7 Bantu populations (N=20), namely North-East Bantus from Kenya, South-West Bantus from Ovambo and Herero, South-East Bantus from Pedi, Southern Sotho, Tswana and Zulu available in the Human Genome Diversity Project (HGDP) (Li, et al. 2008). Overall we evaluated a total of 99,277 single nucleotide polymorphisms (SNPs) that are common to all three datasets under assessment. We mapped the biogeographical origin of the Siddi populations by applying an admixture based method, Geographical Population Structure (GPS) (Elhaik, et al. 2014) that has been successfully employed to trace the accurate origin of various modern populations (Das, et al. 2016; Marshall, et al. 2016). This approach relies on extrapolating the genomic similarity between the query and reference populations to infer the potential biogeographical affiliation of the former using the geographic locations corresponding to the latter. Finally we evaluated our findings in the context of existing historical and anthropological evidences to dissect the genetic relatedness between populations, the fine-scale population structure and recapitulate the population history of the Siddi tribe.

## Material and methods

### Datasets

The datasets used in the present study were obtained from 1000 Genomes project, phase 3 (Genomes Project, et al. 2015), the Human Genome Diversity Project (HGDP) dataset 2 from Stanford University (Li, et al. 2008) and the Reich Lab, Department of Genetics, Harvard Medical School, USA (Auton, et al. 2009; Moorjani, et al. 2013). File conversions and manipulations were performed using EIG v4.2 (Price, et al. 2006), VCF tools (Danecek, et al. 2011) and PLINK (http://pngu.mgh.harvard.edu/purcell/plink/) (Purcell, et al. 2007). All three datasets obtained from 1000 Genomes project, HGDP and the Reich lab were made compatible with each other and merged together using PLINK.

### Population clustering and admixture analysis

Population stratification was estimated using the – – cluster command in PLINK. Multidimensional scaling analysis was performed in PLINK using – – mds command alongside the – – read-genome flag. The multidimensional (mds) plot was generated using R package v3.2.3, and it comprised of genome data pertaining to 504 Africans from 1000 Genomes project, 20 Bantu individuals from HGDP and 14 Siddis from the Reich Lab dataset. The ancestry of all 538 genomes was estimated using ADMIXTURE v1.3 (Alexander, et al. 2009). The ancient splits and genetic relatedness between the populations was inferred using TreeMix v1.13 (Pickrell and Pritchard 2012).

### Tracing the biogeographical origin of Siddis

Biogeographical analysis was performed using the Geographic Population Structure (GPS) algorithm, a highly accurate tool for biogeography estimation that has been demonstrated to effectively trace the origins of several modern populations (Das, et al. 2016; Elhaik, et al. 2014; Marshall, et al. 2016). Given a sample of unknown geographic origin and admixture proportions that correspond to putative ancestral populations, the GPS tool converts the genetic distances between the test sample and the nearest ancient reference populations into geographic distances. The coordinates predicted through GPS should be considered as the last place where the admixture had occurred or the ‘geographical origin’. For individuals of mixed ancestries, GPS coordinates represent the mean geographic locations of their immediate parental populations. All supervised admixture proportions were calculated as illustrated in Elhaik *et al.* (2014) (Elhaik, et al. 2014). In the present study we used the African populations from the 1000 Genomes project and HGDP as reference and interpreted their admixture fractions and geographic locations (latitudnal and longitudnal coordinates) to determine the biogeographical ancestry of the Siddi people.

### Determination of the time of admixture

The level and time of admixture events were estimated using ALDER v.1.02 (Loh, et al. 2013) using a generation time of 25 years. The input files (‘EIGENSTRAT’ format) for ALDER analysis were generated using EIG v. 4.2 (Price, et al. 2006). The Siddi samples were used as the ‘admixpop’ (admixed population) and the African populations from 1000 Genomes project and HGDP were used as the ‘refpops’ (reference populations) during the ALDER analysis.

## Results

### Clustering of populations

The multidimensional (MDS) plot revealed 5 distinct clusters, each corresponding to the African populations derived from the 1000 Genomes project (YRI, GWD, MSL, ESN and LWK), a separate somewhat overlapping cluster of the 7 Bantu populations derived from HGDP (North-East Bantus from Kenya, South-West Bantus from Ovambo and Herero, South-East Bantus from Pedi, Southern Sotho, Tswana and Zulu) that consistent with expectation appeared to be genetically closest to the LWK group and an isolated cluster of the Siddis (fig. 1). While the Siddis emerged as significantly distinct from the rest of the African populations, this plot does provide a deeper insight into the Siddi demographic history revealing them to be genomically closest to LWK and North-East Bantus from Kenya. The Siddi genomes from the Indian state of Karnataka comprised of two groups, Karnataka-1 (N=4) and Karnataka-2 (N=2). We note that consistent with previous findings (Shah, et al. 2011) our MDS plot also revealed the Karnataka-2 Siddis to be genetically closer to the Gujarati Siddis than the other Karnataka-1 group.

**Fig. 1.**
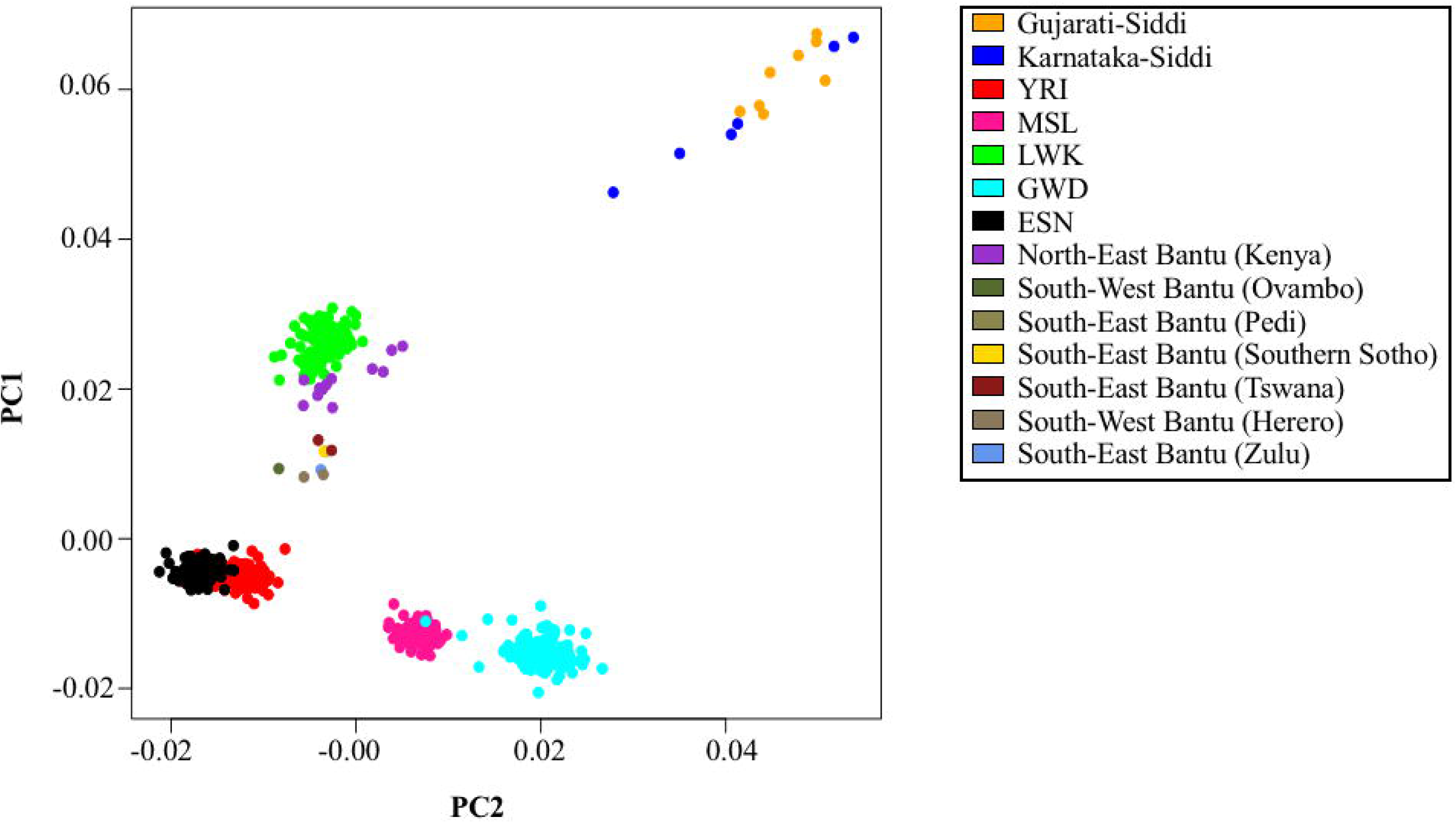
A multidimensional scaling plot of Siddi samples along with African populations from 1000 Genomes project and HGDP. In this scatter plot each point represents an individual. Multidimensional scaling analysis was performed in PLINK and the plot was generated in R v3.2.3. The orange and blue circles designate Siddi populations from Gujarat and Karnataka respectively.

At K=5, the YRI, MSL, LWK, GWD and Siddi genomes were assigned homogeneously to unique populations, although the latter were revealed to share some degree of genetic similarity with the Bantu groups (fig. 2). As may be surmised we found that populations from the same or overlapping geographic location, such as the Nigerians (YRI and ESN) showed a very high degree of genomic similarity. Similar results were obtained for all Bantu groups, including Luhyas (LWK) and North-East Bantus from Kenya, South-West Bantus from Ovambo and Herero, South-East Bantus from Pedi, Southern Sotho, Tswana and Zulu. Siddi genomes were determined to have the highest fraction of component k5 that is unique to them, followed by Bantu (k4), Nigerian (k1), and Mende (k2) ancestral components.

**Fig. 2.**
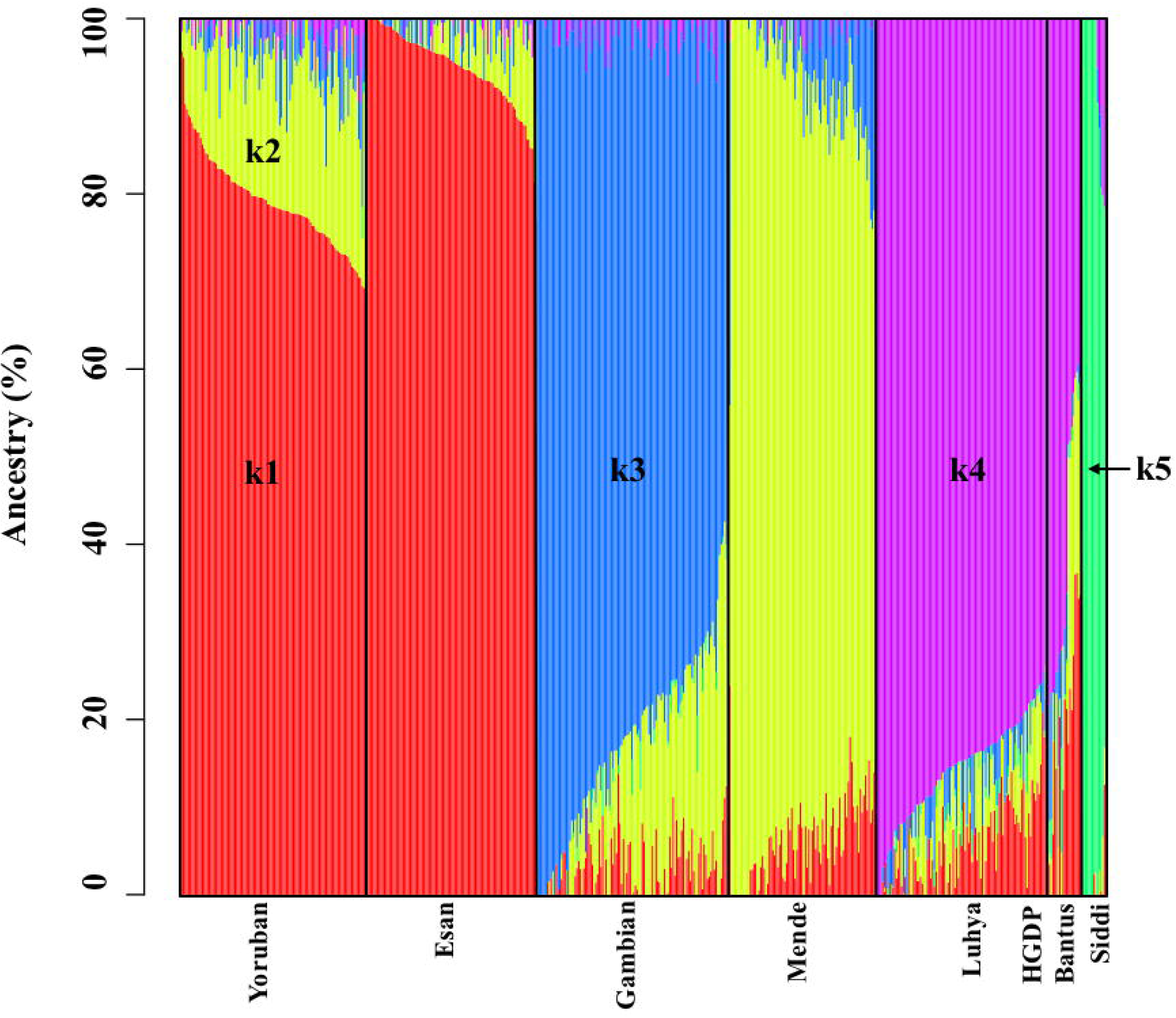
An admixture plot showing the ancestry components of the Indian Siddis along with the African populations. Percent ancestry is plotted on the Y axis. The ancestral components in evaluated genomes was estimated using ADMIXTURE v1.3. To note k1, k2, k3, k4, and k5 represent putative ancestral Nigerian, Mende, Gambian, Bantu, and uniquely Siddi components respectively.

We applied the TreeMix v1.13 model (Pickrell and Pritchard 2012) to investigate the pattern of population splits and mixtures amongst the Indian Siddis and the African populations from the 1000 Genomes project and HGDP. Our findings confirmed the high degree of genetic relatedness between the Siddis and other Bantu populations, especially the Luhyas (LWK) (fig. 3). It further revealed that the African ancestors of Siddis potentially split from the LWK group, ~2700 YBP.

**Fig. 3.**
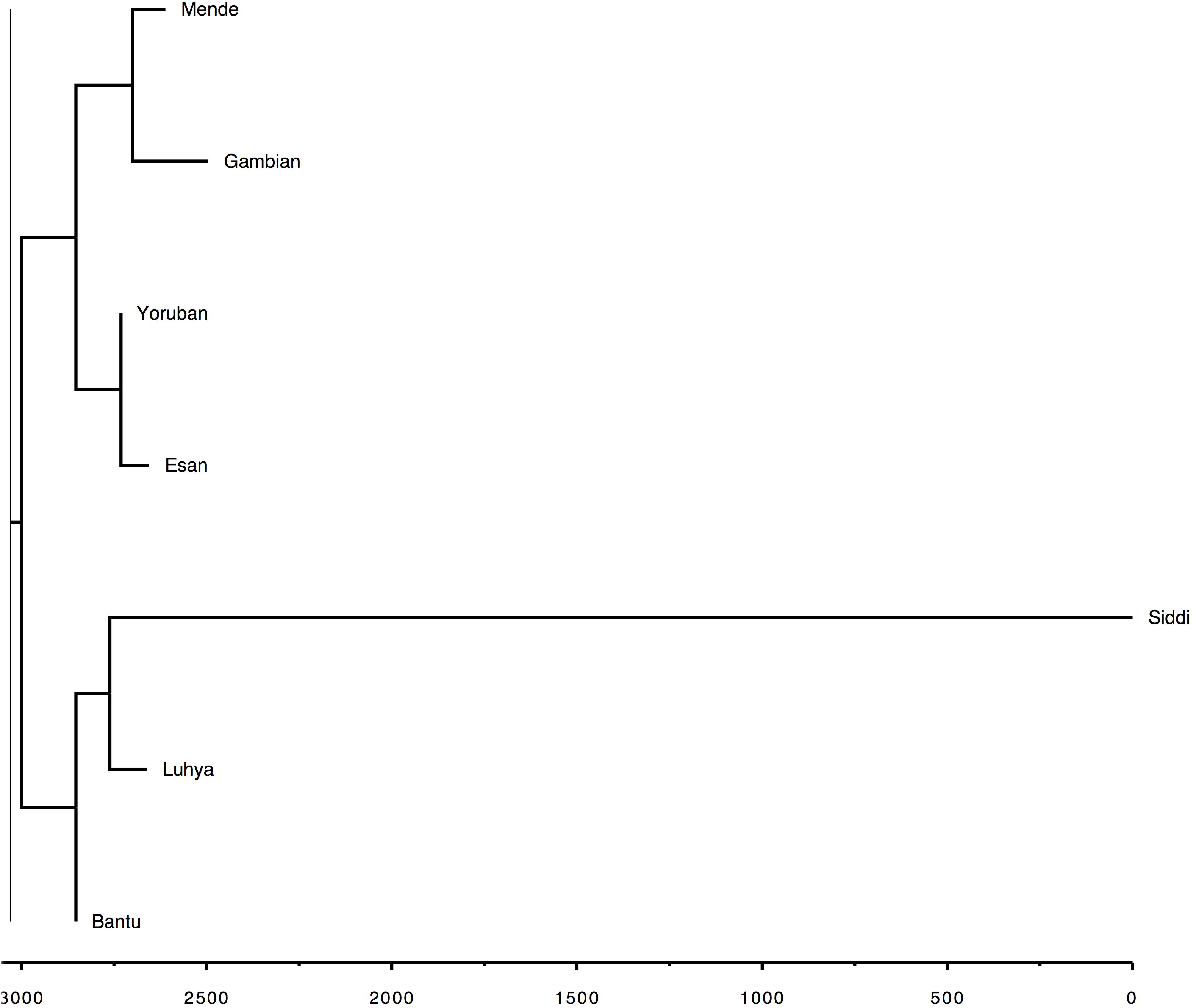
A population tree depicting the relationships between Indian Siddis and the African populations. The population tree was constructed using TreeMix v1.13. The scale bar depicts the time-line in years before present (YBP). This confirmed the high genetic relatedness between Siddis and the Bantu populations, especially Luhyas (LWK), revealing that the Siddis potentially split from them ~2700 YBP.

### Tracing the origin of Siddi populations

All biogeographical inferences were estimated using the GPS algorithm (Elhaik, et al. 2014). GPS assigned 10 out of 14 Siddi samples to west Zambia, while the remaining were positioned at the present day border between Zimbabwe and northeastern Botswana (fig.4). It is noteworthy that the GPS tool assigned the Siddi genomes along the postulated secondary expansion route of the Bantu tribes from the east African Bantu center in the inter-lacustrine region that thrived ~1700 YBP (Pereira, et al. 2001; Phillipson 1993). Strikingly the geographic coordinates of the Siddis as predicted by GPS overlapped with that of likely Bantu settlements in southeastern Africa, from where ancestral African Siddi groups were purportedly transported to India during the 13^th^-19^th^ centuries (Gauniyal, et al. 2008; Nevet 1981; Pereira, et al. 2001; Salas, et al. 2002; Salas, et al. 2004; Shah, et al. 2011).

**Fig. 4.**
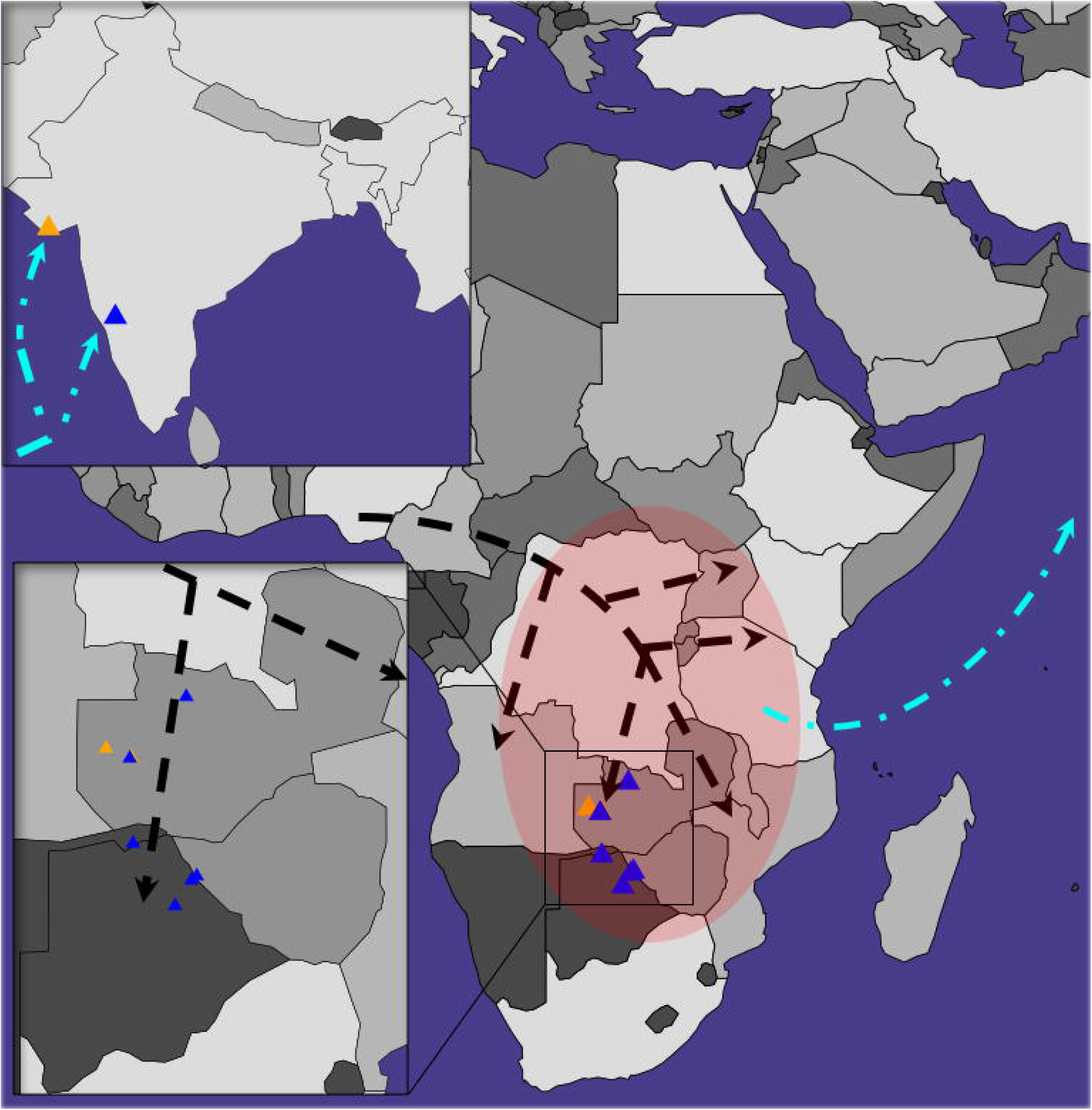
A map depicting the GPS assigned locations of the Indian Siddi genomes. The blue and orange triangles depict Siddi populations from Karnataka and Gujarat respectively. The potential Bantu expansion route through south and southeast Africa is depicted by broken black lines (not to scale). The red oval depicts the secondary Bantu settlements in south and southeast Africa from where ancestral Siddi individuals were likely transported to India during the 13^th^-19^th^ centuries (depicted by broken cyan lines).

### Determination of the date of admixture

To quantitatively estimate the date for the admixture between ancestral Siddis and South Asian populations we applied ALDER v.1.02 (Loh, et al. 2013) that is more precise in comparison with the previously utilized ROLLOFF method (Shah, et al. 2011) and traced back the admixture event to have occurred ~300 YBP or 12.66 generations ago, assuming a generation time of 25 years. Our estimation is in close correspondence with ethnohistorical evidences of ancestral Bantu speaking Siddi groups being imported to India as slaves during the 16^th^ and 17^th^ centuries, by the Portuguese (Bhattacharya 1970; Nevet 1981).

## Discussion

The Siddis are a unique tribal group settled in India whose ancestry is composed of African, South Asian and European components (Bhattacharya 1970; Gauniyal, et al. 2011; Gauniyal, et al. 2008; Ramana, et al. 2001; Shah, et al. 2011; Thangaraj, et al. 1999). Several genetic studies have suggested that they are most closely related to Africans (Gauniyal, et al. 2011; Gauniyal, et al. 2008) and have traced their ancestry to Bantu language speakers from Sub-Saharan Africa (Shah, et al. 2011). The Bantu populations refer to 300-600 African ethnic groups, who speak Bantu languages belonging to the Bantoid subgroup of Benue-Congo branch in the Niger-Congo language family and predominantly occupy central, southeastern and southern Africa (Butt 2006; Nurse 2006). The Bantu expansion is regarded as a crucial demographic event in the history of Sub-Saharan Africa and coincided with the spread of agriculture and iron metallurgy to southern and central regions of the continent (Diamond and Bellwood 2003; Newman 1995; Phillipson 1993). Linguistic, archaeological and genetic evidences are consistent in suggesting that Bantu groups originated in the vicinity of the Cross River valley near the present-day border between Nigeria and Cameroon and their expansion occurred in multiple migratory waves, ~5000 YBP (Greenberg 1964; Huffman 1982; Pereira, et al. 2001; Phillipson 1993; Salas, et al. 2002). A southwest migration led to the dispersal of the proto-Bantu people to the Congo rainforest, while another concomitant southward migration likely progressed along the Atlantic coast, ~3500 YBP. The eastward wave of Bantu dispersal brought them to the inter-lacustrine region around the Great Lakes of east Africa, in southern Uganda and western Tanzania, ~3000 YBP. A subsequent wave of Bantu dispersal initiated from the eastern inter-lacustrine Bantu center, ~1700 YBP southwards to eastern Zimbabwe and eventually Mozambique (Newman 1995; Phillipson 1993). The migratory routes of Bantu populations seemingly converged and overlapped during distinct periods of time leading to the spread of Bantu genomes and languages to a great majority in sub-equatorial Africa (Plaza, et al. 2004). In the present study, we traced the biogeographical origin of the Indian Siddi groups using the GPS tool that mapped them to west Zambia and the present-day border between Zimbabwe and northeastern Botswana, overlapping with one of the principal areas of secondary Bantu settlement in Africa (fig. 4).

Of the 8 Bantu populations included in our analyses, the Indian Siddis were revealed to share the greatest genomic proximity with the Luhyas from Webuye (LWK) and North-East Bantu populations of Kenya (fig. 1 and fig. 2). This was further supported by the results of our population tree analysis applying the TreeMix v1.13 algorithm that also indicated the Siddis to be genetically close to the Bantu populations, in particular the Luhyas and likely having split from them ~2700 YBP (fig. 3). Notably while previous studies discerned a predominant African component in the Siddi ancestry, their genetic relatedness when compared to East (LWK) and West (YRI) African genomes had remained unresolved (Shah, et al. 2011). Our findings are also concordant with a secondary Bantu dispersal route, southwards from their east African center along the shores of Lake Malawi, via eastern Zimbabwe and along the Ruvuma river that forms the border between Tanzania and Mozambique (Pereira, et al. 2001) that likely brought ancestral Siddi groups to settlement sites in southeast Africa from where they were disseminated to India.

Historically southeast Africa had emerged as an important source of slaves from the 16^th^ century onwards, when individuals from Mozambique and Madagascar constituted a major proportion of the slaves being shipped by the Portuguese, primarily to former European colonies, such as the Caribbean and Brazil in the Americas (Thomas 1998). Several lines of evidences suggest a concordant time-line, during the 16^th^-19^th^ centuries, when ancestral Siddi people were transported by the Portuguese from Mozambique and its neighboring areas in southeast Africa, and eventually sold to regional Indian princes and chieftains, to serve as slaves and soldiers (Bhattacharya 1970; Nevet 1981). We sought to delineate the time-scale of the admixture event in the Siddi population history by applying the ALDER algorithm that employs an improved weighted linkage disequilibrium based method to investigate admixture (Loh, et al. 2013). Our analysis dates the admixture between Siddis and South Asians to ~300 YBP or 12.66 ± 2.11 generations ago, using a generation time of 25 years. This estimation is older and likely more precise than an earlier proposed timeline of 200 YBP or 8 ± 1 generations ago (Shah, et al. 2011) that was calculated using the less accurate ROLLOFF algorithm (Loh, et al. 2013). Our proposed time-scale of admixture appears to be congruent with historical evidences, which suggest that a large number of African ancestors of the Siddis were imported to India from Mozambique between 1680 A.D and 1720 A.D (Gauniyal, et al. 2008; Nevet 1981).

To the best of our knowledge this is the first study to investigate the biogeographical origin of the Siddis, a unique tribe of African ancestry from India. Our findings when taken together with historical, anthropological and genetic evidences enable a reconstruction of the distinctive population history of Siddis and offer unequivocal insights into the demographic factors that likely contributed to the contemporary Siddi genomes.

## References

Alexander DH, Novembre J, Lange K 2009. Fast model-based estimation of ancestry in unrelated individuals. Genome Res 19: 1655–1664. doi: 10.1101/gr.094052.109

Auton A, et al. 2009. Global distribution of genomic diversity underscores rich complex history of continental human populations. Genome Res 19: 795–803. doi: 10.1101/gr.088898.108

Barnabas S, Apte RV, Suresh CG 1996. Ancestry and interrelationships of the Indians and their relationship with other world populations: a study based on mitochondrial DNA polymorphisms. Ann Hum Genet 60: 409–422.

Bhattacharya D 1970. Indians of African origin. Cah. Etud. Afr. 10: 579–582.

Butt JJ. 2006. The Greenwood dictionary of world history: Greenwood Publishing Group. D.K B 1969. Anthropometry of a Negroid population in India: Siddis of Gujarat. J. Anthropol. Soc. Nippon 77: 254–256.

Danecek P, et al. 2011. The variant call format and VCFtools. Bioinformatics 27: 2156–2158. doi: 10.1093/bioinformatics/btr330

Das R, Wexler P, Pirooznia M, Elhaik E 2016. Localizing Ashkenazic Jews to Primeval Villages in the Ancient Iranian Lands of Ashkenaz. Genome Biol Evol 8: 1132–1149. doi: 10.1093/gbe/evw046

Diamond J, Bellwood P 2003. Farmers and their languages: the first expansions. Science 300: 597–603. doi: 10.1126/science.1078208

Elhaik E, et al. 2014. Geographic population structure analysis of worldwide human populations infers their biogeographical origins. Nat Commun 5: 3513. doi: 10.1038/ncomms4513

Gauniyal M, Aggarwal A, Kshatriya GK 2011. Genomic structure of the immigrant Siddis of East Africa to southern India: a study of 20 autosomal DNA markers. Biochem Genet 49: 427–442. doi: 10.1007/s10528-011-9419-7

Gauniyal M, Chahal SM, Kshatriya GK 2008. Genetic affinities of the Siddis of South India: an emigrant population of East Africa. Hum Biol 80: 251–270. doi: 10.3378/1534-6617-80.3.251

Genomes Project C, et al. 2015. A global reference for human genetic variation. Nature 526: 68–74. doi: 10.1038/nature15393

Greenberg J. 1964. Historical inferences from linguistic research in Sub-Saharan Africa. In: Butler J, editor. Boston University Papers in African History I. Boston, MA: Boston University Press. p. 1–15.

Huffman TN 1982. Archaeology and the ethnohistory of the African Iron Age. Annu Rev Anthropol 11: 133–150.

Kivisild T, et al. 1999. Deep common ancestry of indian and western-Eurasian mitochondrial DNA lineages. Curr Biol 9: 1331–1334.

Li JZ, et al. 2008. Worldwide human relationships inferred from genome-wide patterns of variation. Science 319: 1100–1104. doi: 10.1126/science.1153717

Lodhi A 1992. African settlements in India. Nordic Journal of African Studies 1: 83–86.

Loh PR, et al. 2013. Inferring admixture histories of human populations using linkage disequilibrium. Genetics 193: 1233–1254. doi: 10.1534/genetics.112.147330

Marshall S, Das R, Pirooznia M, Elhaik E 2016. Reconstructing Druze population history. Scientific Reports 6: 35837. doi: 10.1038/srep35837

Metspalu M, et al. 2004. Most of the extant mtDNA boundaries in south and southwest Asia were likely shaped during the initial settlement of Eurasia by anatomically modern humans. BMC Genet 5: 26. doi: 10.1186/1471-2156-5-26

Misra VN 2001. Prehistoric human colonization of India. J Biosci 26: 491–531.

Moorjani P, et al. 2013. Genetic evidence for recent population mixture in India. Am J Hum Genet 93: 422–438. doi: 10.1016/j.ajhg.2013.07.006

Narang A, et al. 2010. IGVBrowser--a genomic variation resource from diverse Indian populations. Database (Oxford) 2010: baq022. doi: 10.1093/database/baq022

Nevet A. 1981. John the Britto. Bangalore, India: Loyal Mandira.

Newman JL. 1995. The peopling of Africa: a geographic interpretation: Yale University Press.

Nurse D. 2006. Bantu Languages. In: Bown K, editor. Encyclopedia of Language and Linguistics. Amsterdam: Elsevier.

Pereira L, et al. 2001. Prehistoric and historic traces in the mtDNA of Mozambique: insights into the Bantu expansions and the slave trade. Ann Hum Genet 65: 439–458. doi: 10.1017/S0003480001008855

Phillipson DW. 1993. African archaeology. Cambridge: Cambridge University Press.

Pickrell JK, Pritchard JK 2012. Inference of population splits and mixtures from genome-wide allele frequency data. PLoS Genet 8: e1002967. doi: 10.1371/journal.pgen.1002967

Plaza S, et al. 2004. Insights into the western Bantu dispersal: mtDNA lineage analysis in Angola. Hum Genet 115: 439–447. doi: 10.1007/s00439-004-1164-0

Price AL, et al. 2006. Principal components analysis corrects for stratification in genome-wide association studies. Nat Genet 38: 904–909. doi: 10.1038/ng1847

Purcell S, et al. 2007. PLINK: a tool set for whole-genome association and population-based linkage analyses. Am J Hum Genet 81: 559–575. doi: 10.1086/519795

Ramana GV, et al. 2001. Y-chromosome SNP haplotypes suggest evidence of gene flow among caste, tribe, and the migrant Siddi populations of Andhra Pradesh, South India. Eur J Hum Genet 9: 695–700. doi: 10.1038/sj.ejhg.5200708

Ratnagar S. 1995. Archaeological perspectives of early Indian societies. In: Thapar R, editor. Recent perspectives of early Indian history. Mumbai, India: Popular Prakashan. p. 1–52.

Salas A, et al. 2002. The making of the African mtDNA landscape. Am J Hum Genet 71: 1082–1111. doi: 10.1086/344348

Salas A, et al. 2004. The African diaspora: mitochondrial DNA and the Atlantic slave trade. Am J Hum Genet 74: 454–465. doi: 10.1086/382194

Shah AM, et al. 2011. Indian Siddis: African descendants with Indian admixture. Am J Hum Genet 89: 154–161. doi: 10.1016/j.ajhg.2011.05.030

Tamang R, Singh L, Thangaraj K 2012. Complex genetic origin of Indian populations and its implications. J Biosci 37: 911–919.

Thangaraj K, Ramana GV, Singh L 1999. Y-chromosome and mitochondrial DNA polymorphisms in Indian populations. Electrophoresis 20: 1743–1747.

Thomas H. 1998. The slave trade - the history of the Atlantic slave trade. London: Macmillan Publishers Ltd.

